# Biosensor capability of the endometrium is mediated in part, by altered miRNA cargo from conceptus-derived extracellular vesicles

**DOI:** 10.1101/2023.10.27.564369

**Authors:** Tiago H. C. De Bem, Alessandra Bridi, Haidee Tinning, Rafael Vilar Sampaio, Irene Malo-Estepa, Dapeng Wang, Elton J. R. Vasconcelos, Ricardo Perecin Nociti, Ana C. F. C. M. de Ávila, Juliano Rodrigues Sangalli, Igor Garcia Motta, Gilmar Arantes Ataíde, Júlio C. B. da Silva, Yeda Fumie Watanabe, Angela Gonella-Diaza, Juliano C. da Silveira, Guilherme Pugliesi, Flávio Vieira Meirelles, Niamh Forde

## Abstract

We tested the hypothesis that the biosensor capability of the endometrium is mediated in part, by the effect of different cargo contained in the extracellular vesicles secreted by the conceptus during the peri-implantation period of pregnancy. We transferred *Bos taurus taurus* embryos of different origin: *In vivo* (high developmental potential (IV)), *in vitro* (intermediate developmental potential (IVF)), or cloned (low developmental potential (NT)), into *Bos taurus indicus* recipients. Extracellular vesicles (EVs) recovered from Day 16 conceptus conditioned medium were characterized and their microRNA (miRNA) cargo sequenced alongside RNA sequencing of their respective endometria. There were substantial differences in the endometrial response to *in vivo* Vs *in vitro* and *in vivo* Vs cloned conceptuses (1153 and 334DEGs respectively) with limited differences between *in vitro* Vs cloned conceptuses (36 DEGs). miRNA cargo was similar between all three groups (426 common cargo) differences between *in vivo* and cloned (8 miRNAs), and *in vivo* and *in vitro* (6 miRNAs) observed. Treatment of endometrial epithelial cells with mimic or inhibitors for miR-128 and miR-1298 changes to the proteomic content of target cells (96, and 85 respectively) of which mRNAs are altered in the endometrium *in vivo* (*PLXDC2, COPG1, HSPA12A, MCM5, TBL1XR1, and TTF*). In conclusion, we have determined that the biosensor capability of the endometrium is mediated in part, by its response to different EVs miRNA cargo produced by the conceptus during the peri-implantation period of pregnancy.

**SIGNIFICANCE STATEMENT:** During the peri-implantation period of pregnancy in mammals, the endometrium acts as a biosensor for the developmental competency of the embryo. However, the mechanism by which biosensor capability of the endometrium is established, remains elusive. In this study, we show that embryos of different developmental competencies have distinct microRNA cargo contained in their extracellular vesicles (EVs). Exposure of the endometrium to these conceptuses alters the transcriptional response of the endometrium during the process of pregnancy recognition. This differential response is mediated in part, by the delivery and action of the these differentially abundant microRNAs into EVs. Here we propose differences in EV-mediated miRNA cargo are responsible in part for this biosensor capability of the endometrium.

## INTRODUCTION

In placental mammals, the majority of reproductive wastage occurs in the first 2-3 weeks of pregnancy with a specific wave of loss occurring during the peri-implantation period (1). Establishing successful early pregnancy requires communication between the gametes and the maternal environment as well as successful bi-directional communication between the developing conceptus (embryo and extra-embryonic membranes) and the uterine environment. Irrespective of the species studied, or indeed the type of conceptus present, there is an optimum window of uterine receptivity (UR) to implantation (2–4). Furthermore, recent data in multiple species indicates that the endometrium is capable of ‘sensing’ the type of embryo present (5–7). Studies in mammals with diverse implantation strategies (invasive implantation in humans and superficial implantation in cattle) have shown that embryos with different developmental competencies i.e. embryos more likely to give rise to successful pregnancy (*in vivo* produced) versus those that do not (*in vitro*, clones), are sensed by the endometrium during pregnancy recognition/implantation and the endometrium modifies its transcriptome accordingly (5–7). Not only does the endometrium establish UR to implantation but it also modifies this receptivity depending on the quality of embryo present. However, the mechanism by which the endometrium does this still remains elusive.

In cattle, coordinate with conceptus elongation, the trophectoderm cells produce and secrete increasing concentrations of the pregnancy recognition signal IFNT (8). In addition to this secretion of IFNT, the conceptus produces additional proteins (4, 9–11) that induce transcriptional changes in the endometrium. Interestingly, data from Bauersachs et al., (6) demonstrates that exposure of the endometrium to the conceptus induced a larger transcriptional response than exposure to a type 1 Interferon response alone *in vivo* i.e. factors other than IFNT alter the endometrial transcriptome. In work by Bauersachs (12) no changes in IFNT production by embryos with different developmental competencies was determined and, therefore, the biosensor capability of the endometrium is not mediated only by production of different concentrations of IFNT and subsequent actions on the endometrium.

Recently, extracellular vesicles (EVs) have emerged as a non-traditional form of cell-to-cell communication. EVs are membrane-bound vesicles (classified depending on their size and biogenesis) and contain lipids, proteins, as well as RNA (both coding and non-coding RNAs) molecules that are capable of being incorporated into target tissues (13). Data in mouse show that EVs produced by embryonic stem cells (ESC) are delivered to the trophectoderm lineage to enhance embryo implantation in the endometrium (14). One of the first reports of this phenomenon in the uterus was through the incorporation of endogenous retroviral envelope proteins that are shed from the endometrial epithelium and incorporated into the trophectoderm cells of sheep conceptuses with more recent data confirming that the mode of transfer of these retroviral particles was via EVs (specifically exosomes; (15, 16)). This phenomenon has been reported in other species e.g. sheep and cattle at different points on the reproductive axis (17). More recently, it has been demonstrated that miRNAs from the embryo are detectable in culture media, although it is not clear if these are EV-derived (18, 19). Additionally, bovine blastocysts produced *in vivo* or *in vitro* are capable to secrete different amounts of extracellular vesicles containing different miRNA contents (20).

The **<u>overarching hypothesis</u>** tested is that the biosensor capability of the endometrium is mediated in part, by loading of different EV cargo derived from the conceptus during the pregnancy recognition period. To test this hypothesis, we produced embryos from *Bos taurus taurus* individuals with different developmental potential and transferred them into *Bos taurus indicius* recipients to determine how the biosensor capability of the endometrium is mediated.

## RESULTS

### Animal Model and EV characterization

Following production of *in vivo*, *in vitro*, and cloned *Bos taurus taurus* embryos, blastocysts were transferred into synchronized *Bos taurus indicus* recipients (Figure 1). Fibroblasts from both cell linages (male and female) were evaluated by flow cytometry with 79.6% of cells positive for GFP and 64.4% of cells positive for GFP in the female line (Supplementary Figure 1). Concentrations of plasma progesterone (P4) in recipients at the day of embryo transfer (day 7) or at slaughter (day 16) did not differ (P>0.1) among groups (Figure 2A) nor between day 7 and day 16. On Day 16, 28 conceptuses were recovered from recipients. Sexing of the conceptuses revealed six *in vivo*-derived conceptuses (IV: 3 females, 3 males), 10 *in vitro*-derived (IVF: 7 females, 3 males), and 12 cloned conceptuses (NT: 5 females, 7 males; Supplementary Figure 2). In cloned embryos, green fluorescent protein (GFP) was detected in fibroblast cells (Figure 2B), blastocysts prior to transfer (Figure 2C), and recovered conceptuses (Figure 2D), fibroblast and conceptuses (Figure 2E) as well as adjacent to the LE of the endometrium on Day 16.

**Figure 1.**
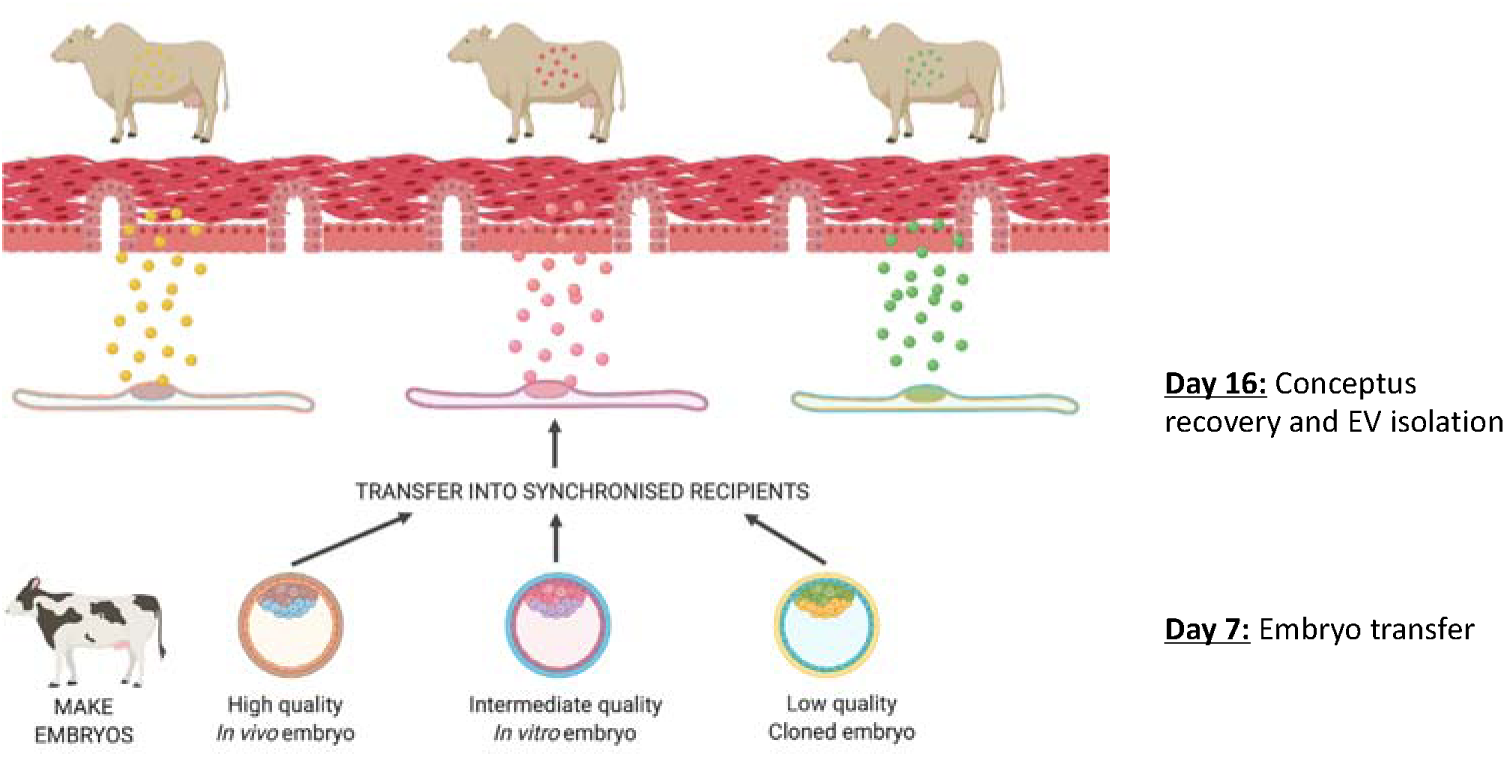
Schematic diagram of animal model. We tested the hypothesis that the biosensor capability of the endometrium is mediated in part by different cargo present in extracellular vesicles produced by the conceptus during the pregnancy recognition period. To test this hypothesis *Bos taurus taurus* embryos were produced to generate both male and female high quality *in vivo* embryos (gold), intermediate quality *in vitro* embryos (pink), or low-quality cloned embryos (green). Embryos were transferred into synchronised *Bos taurus indicus* recipients and conceptuses and endometrial tissue were recovered on Day 16, cultured for 8 h and extracellular vesicles (EVs) characterised and cargo examined.

**Figure 2.**
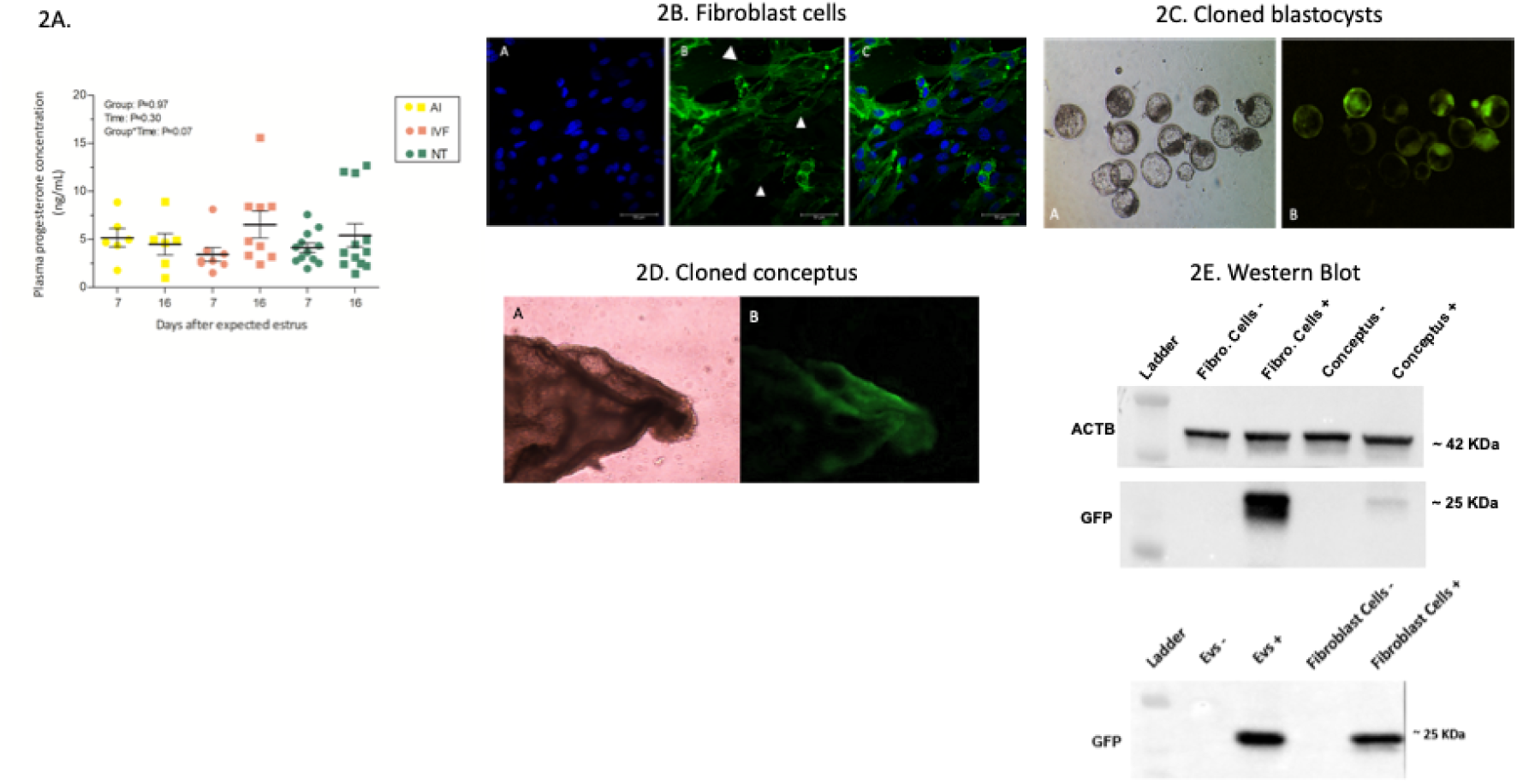
Animal model validation. **(A)** Plasma progesterone concentrations measured on Day 7 (day of embryo transfer) and Day 16 (day of maternal recognition of pregnancy) in recipient animals where *in vivo* (yellow), *in vitro* (pink), or cloned (green) embryos were transferred. **(B)** Representative image of GFP present in fibroblast cells used to generated cloned embryos. GFP expressing **(C)** blastocysts and **(D)** day 16 conceptuses. **(E)** Western blot probing for GFP in fibroblast cells and conceptuses, **(F)** GFP-detected adjacent to luminal epithelium (LE) endometrial on Day 16.

Analyses of EVs derived from conceptus conditioned medium (CCM) revealed no difference in particle concentration (mean concentration 2.15 x 10^9^ particles/mL; *in vivo*: 1.11 x 10^9^, *in vitro*: 2.14 x 10^9^, and clones; 2.69 x 10^9^ particles/mL, Figure 3A), nor size (*in vivo*: 155.35nm (±12.86nm), *in vitro*: 159.14nm (±13.55nm), and clones; 171.2nm (±23.45nm) (Figure 3B). Western blot analysis of markers of EVs detected ALIX in our isolated EVs but the absence of GRP78 (Figure 3C), while TEM demonstrated classical EV shape (Figure 3D+E).

**Figure 3.**
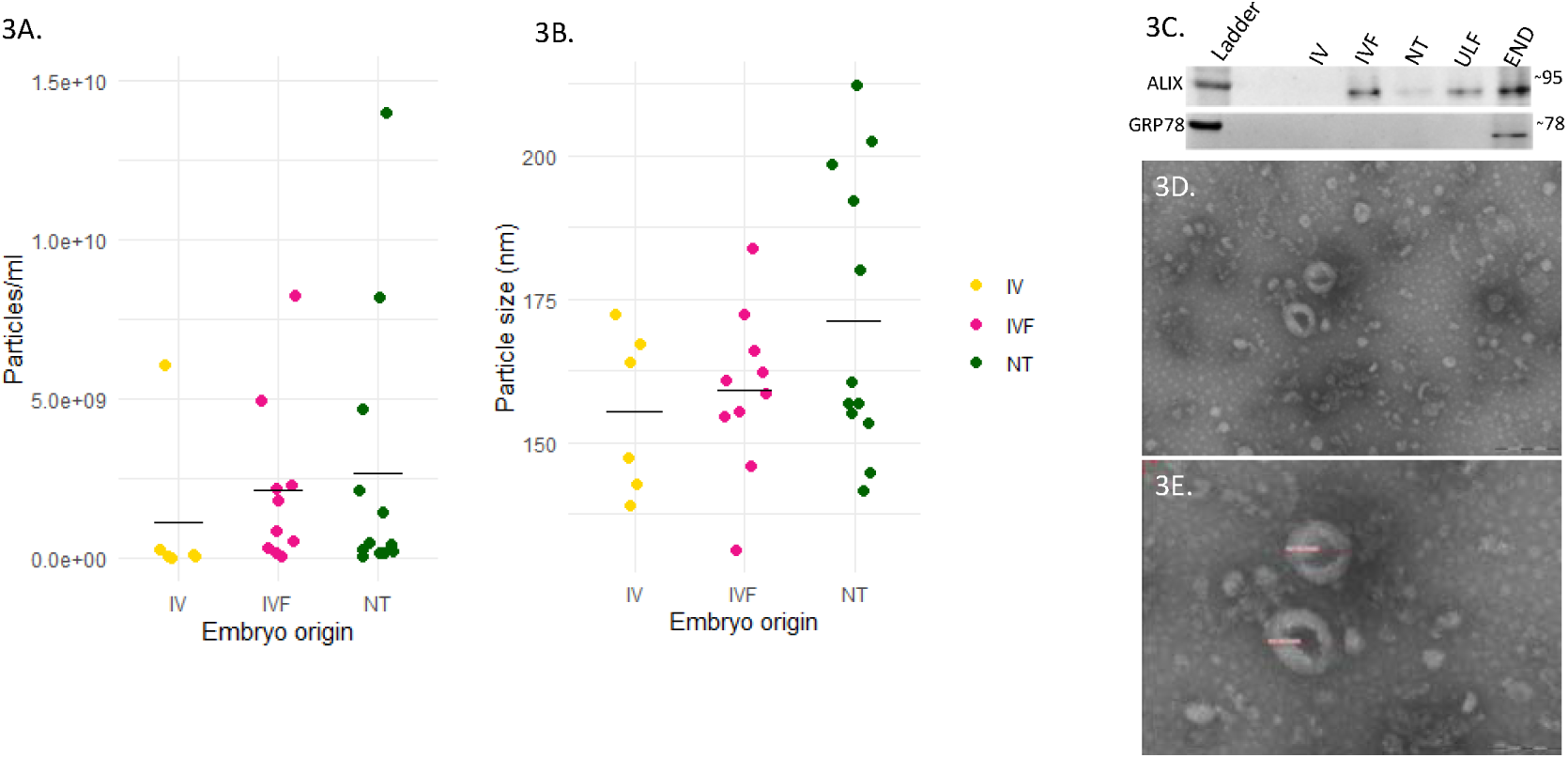
Characterization of EVs derived from conceptuses. **(A+B)** Nano-particle tracking analysis of EVs derived from Day 16 CCM from *in vivo* (yellow), *in vitro* (pink) or cloned (green) conceptuses. No difference in **(A)** particle number concentration or **(B)** size of EVs were detected. **(C)** Western blot analysis of EV markers of EVs ALIX and GRP78 in EVs from IVF, NT, ULF, and endometrium. **(D+E)** TEM of EVs from conceptuses demonstrating the expected size and classical EV shape.

### The endometrium responds differently to conceptuses with different developmental potential and sex

To determine if the endometrium responds to conceptuses with different developmental competencies, we performed RNAseq analysis on the intercaruncular endometrium of recipients on Day 16 of pregnancy. In total 18,077 transcripts containing more than >200 nucleotides with a count >5 were detected in the endometrium. Heatmap analysis demonstrated the overall transcriptional profile of endometrium clustered according to developmental competency of the conceptus to which it was exposed (Figure 4A). Endometria exposed to *in vivo* conceptuses compared to *in vitro* produced conceptuses on Day 16 displayed altered expression of 1153 transcripts (differentially expressed genes – DEGs: 926 of which were upregulated, 227 down regulated: Supplementary Table 1). Overrepresented pathways associated with all 1153 DEGs i.e. irrespective of the direction of change, are involved in molecular functions of integrin binding and cell adhesion molecule binding (Supplementary Table 2), while overrepresented pathways included MAPK signaling pathway, Focal adhesion, and Wnt signaling pathway (Supplementary Table 3). Of the up-regulated transcripts, Oxytocin signaling pathway, hedgehog signaling, and ECM-receptor interactions were overrepresented (Supplementary Table 4), as well as the molecular function ontology of integrin binding (Supplementary Table 5). In contrast, the down-regulated genes (*in vivo* V *in vitro*) were associated with biological processes of positive regulation of secretion by cell, secretory pathway, and positive regulation of exocytosis (Supplementary Table 6) with genes overrepresented in pathways involved in MAPK signaling pathway, Oxytocin signaling pathway, Ras signaling pathway, and Focal adhesion (Supplementary Table 7).

**Figure 4.**
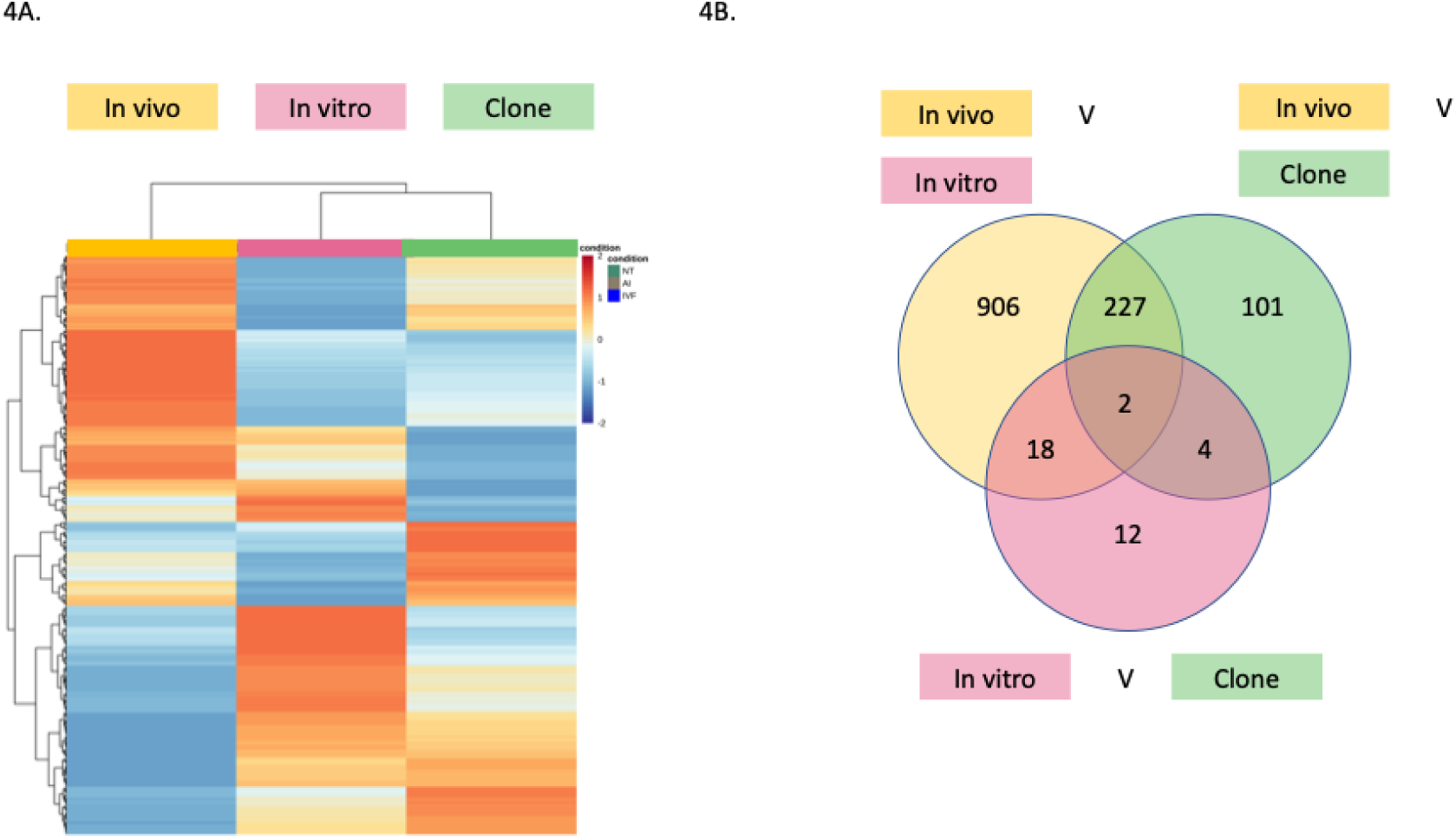
Endometrial response to conceptuses with different developmental competencies. **(A)** Heatmap analyses of transcriptomic response (measured by bulk RNA sequencing) of the endometrium on Day 16 of confirmed pregnancy following transfer of embryos with different developmental competencies on Day 7 to synchronised recipients. Specific clustering of the transcriptional response is observed for endometria exposed to high-quality *in vivo* produced embryos (LHS, gold colour (n=6), intermediate-quality *in vitro* embryos (middle, pink colour (n=10), or low-quality embryos (RHS, green colour (n=10) confirming the biosensor capability of the endometrium previously proposed by (5, 6, 34) (B) Venn diagram with the numbers of differentially expressed genes identified between endometria exposed to embryos with different developmental competencies (IV, IVF and NT).

Exposure of the endometrium to *in vivo* (high-quality) compared to cloned embryos (low developmental potential) altered 334 transcripts of which 291 were upregulated and 43 downregulated (Supplementary Table 8). These were associated with molecular functions of voltage-gated potassium channel complex and potassium channel complex (Supplementary Table 9) and pathways including Axon guidance, Calcium signaling pathway and MAPK signaling pathway (Supplementary Table 10).

In contrast, only 36 DEGs were altered in endometrium exposed to *in vitro* and cloned conceptuses (Supplementary Table 11), all of which were upregulated in *in vitro*-exposed endometrium and are overrepresented in biological process including protein/peptide transport (Supplementary Table 12), and pathways of Drug metabolism - cytochrome P450, and Glutathione metabolism (Supplementary Table 13). Embryos produced by assisted reproductive technologies altered a consistent signature of 227 DEGs (Figure 4B; Supplementary Table 14) compared to *in vivo* produced embryos, but were not associated with any overrepresented categories. While only 2 genes were different in all three groups (Figure 4B. *CHSY3*; chondroitin sulfate synthase 3, and *SLC24A3*; solute carrier family 24 member 3). We also determined that, irrespective of the type of conceptus present the sex of the conceptus altered the transcriptional response independently (Figure 5).

**Figure 5.**
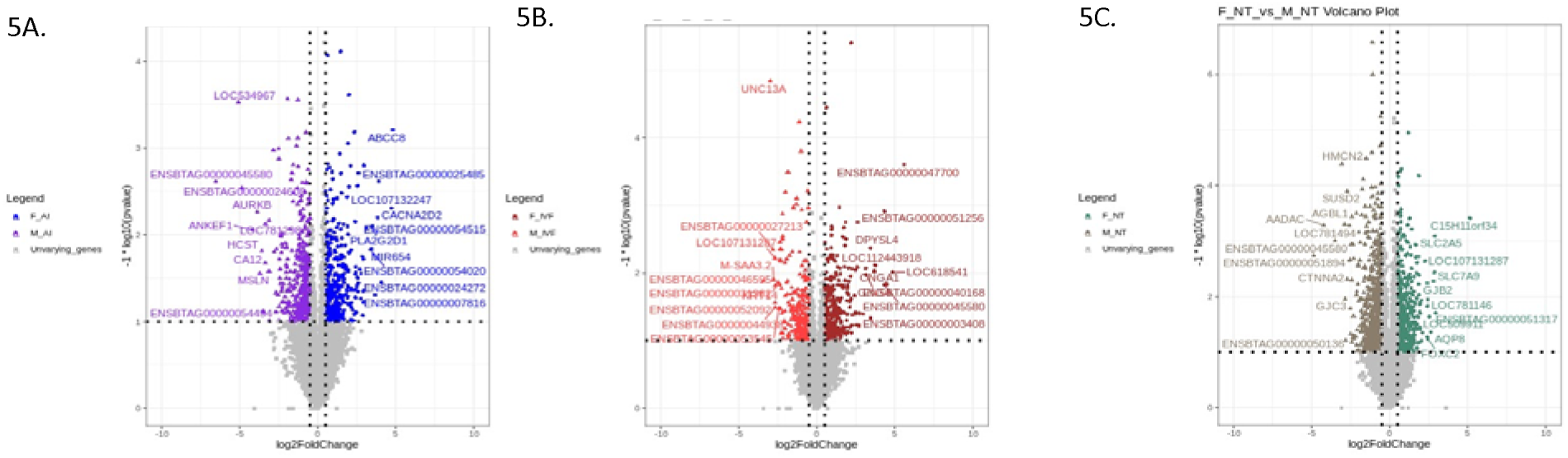
Endometrial response to conceptuses with different developmental competencies. Volcano plots showing differences in transcriptional response of the endometrium to male compared to female conceptuses produced (A) *in vivo*, (B) *in vitro*, or (C) cloned conceptuses on Day 16 of pregnancy.

### Small RNA sequencing of conceptus-derived EVs

To understand if there are differences in miRNA cargo of EVs we sequenced the composition of micro RNAs in EVs. In total, 655 different miRNAs were identified as cargo in day 16 conceptus-derived EVs (Figure 6; Supplementary Table 15). Four hundred and sixty-nine, 540, and 580 miRNAs were identified in EVs derived from *in vivo*, *in vitro*, and cloned conceptuses, respectively. Of these, 426 were common to all the 3 groups and 72 were unique to cloned conceptuses. Cloned conceptuses shared similar miRNA EV cargo with *in vivo* (43 miRNAs) and *in vitro*-produced conceptuses (114 miRNAs) (Figure 6). Filtering of these miRNAs to those with a fold-change >1 or <-1, and p-adjusted value of <0.05 identified no significant differences in miRNA cargo from EVs derived from conceptuses produced *in vitro* or via cloning. In contrast, 6 miRNAs were differentially expressed in EVs derived from *in vivo* compared to *in vitro* conceptuses, 2 of which were downregulated (bta-miR-320a and bta-miR-3432a) and 4 upregulated (bta-miR-1248, bta-miR-2892, bta-miR-2904, and bta-miR-12034). In total, 8 miRNAs were differentially abundant in EVs from *in vivo* compared to cloned conceptuses, 5 downregulated (bta-miR-128, bta-miR-139, bta-miR-218, bta-miR-219-3p, and bta-miR-1298) and 3 upregulated (bta-miR-154c, bta-miR-2419-5p, and bta-miR-2892).

**Figure 6.**
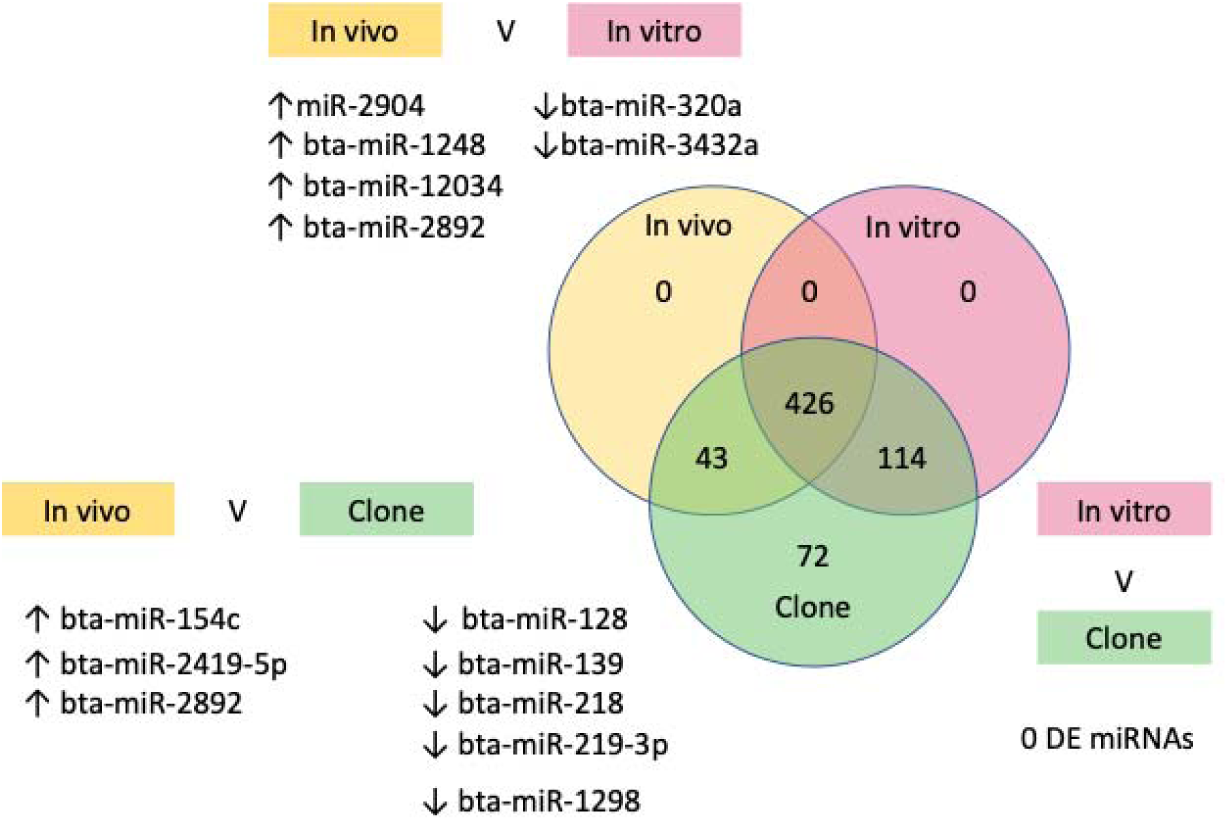
Micro RNA cargo of EVs from Day 16 conceptuses. Venn diagram analyses of miRNA cargo of EVs isolated from CCM. Media was derived from *in vivo*, *in vitro*, or cloned conceptuses cultured *in vitro* for 8 h. EVs were characterised as per ISEV guidelines and miRNA profiling carried out via miRNA sequencing.

### Functional analyses of selected EV-miRNAs

We selected miR-128 and miR-1298 for further analysis as these were EV-derived miRNA with the largest fold change difference between embryos with the most divergent developmental competency i.e. *in vivo* and clones. Following removal of off-target effects, treatment of primary bovine endometrial epithelial cells with miR-128 mimic altered expression of 863 proteins, while treatment with inhibitor altered 562 (Figure 7: Supplementary Table 16). A core of 96 proteins were modified by miR-128 mimic and inhibitor.

**Figure 7.**
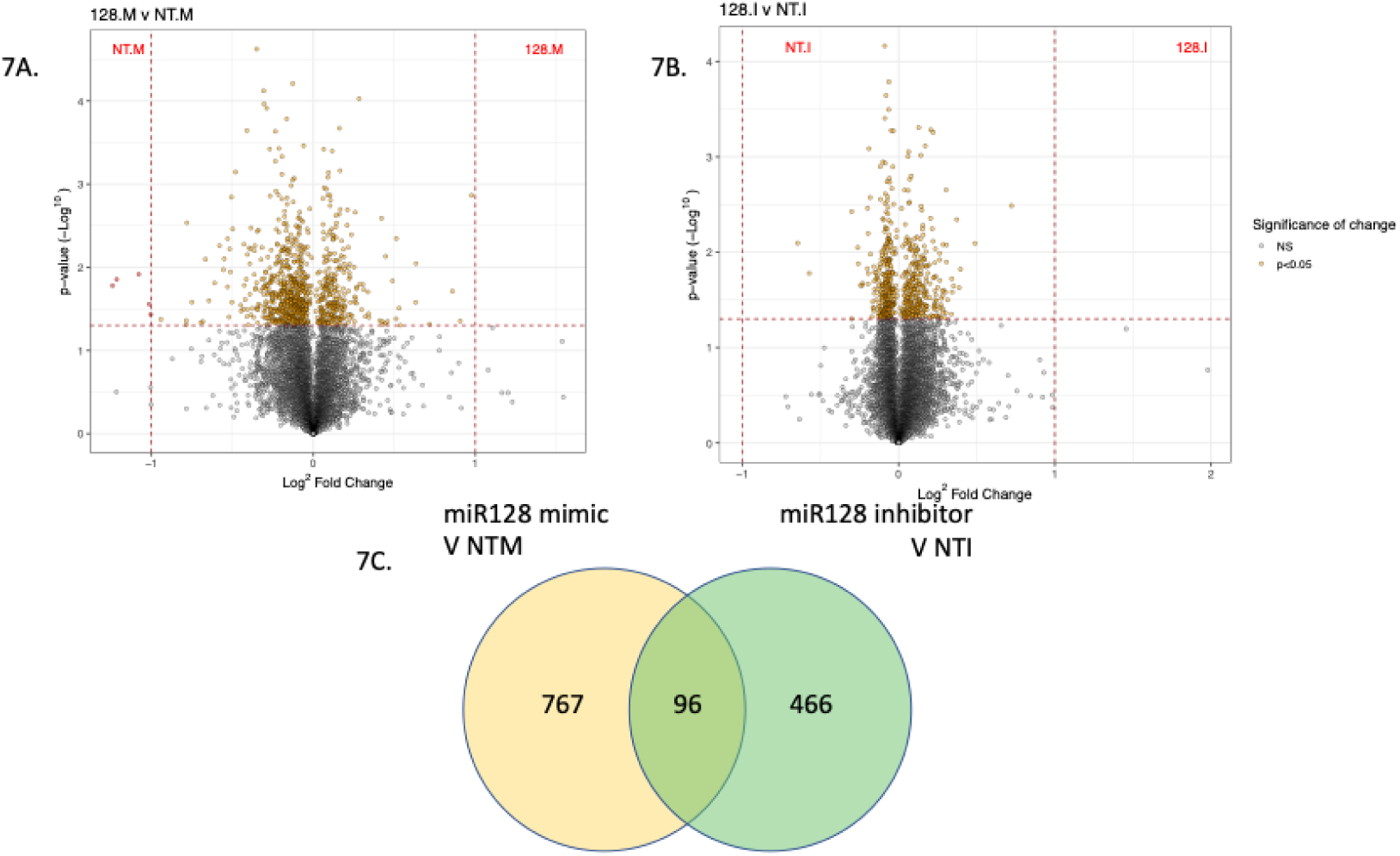
In vitro regulation of proteins by miR128. Volcano plots demonstrating differentially abundant proteomic analysis of bovine endometrial epithelial cells transfected with (A) non-targeting mimic or miR-128 mimic or (B) non-targeting inhibitor or miR-128 inhibitor for 48 h (n=3 biological replicates). Differences were determined by tandem mass tag mass spectrometry analysis of protein lysates from the cells.

Of the proteins altered by miR-128 *in vitro*, the expression of 9 proteins increased, and 2 decreased with their corresponding transcripts are also altered in endometria exposed to cloned *Vs in vivo* conceptuses. One protein (plexin domain containing 2 (PLXDC2) modified by miR-128 mimic and inhibitor *in vitro* and also increased in expression in endometria exposed to *in vivo Vs* cloned conceptuses on Day 16.

For miR-1298, removal of off-target effects revealed 949 proteins modified by mimic treatment alone, while inhibitor treatment altered 562 of which 85 were common to both treatments (Figure 8: Supplementary Table 17). Of these 85 proteins whose abundance was altered by miR-1298, expression of mRNA for 5 of these proteins also changed in the endometrium (COPG1, HSPA12A, MCM5, TBL1XR1, and TTF2). There was minimal overlap in proteins regulated by these miRNAs indicating miRNA specificity in terms of their targets and minimal numbers of these are identified a predicted targets (miRGene DB punch out to Targetscan identified 161 predicted targets for bovine miR-1298).

**Figure 8.**
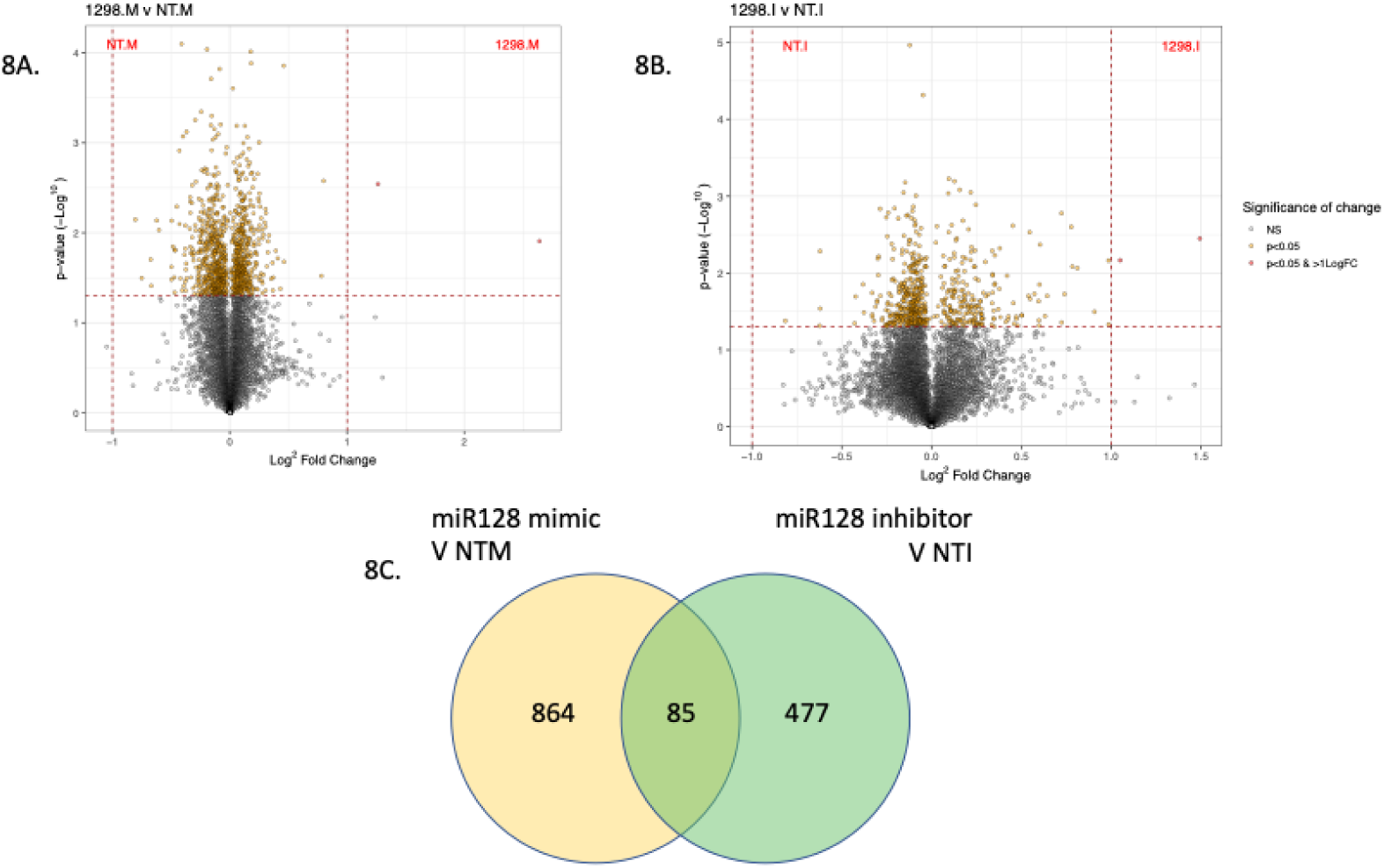
In vitro regulation of proteins by miR1298. Volcano plots demonstrating differentially abundant proteomic analysis of bovine endometrial epithelial cells transfected with (A) non-targeting mimic or miR-1298 mimic or (B) non-targeting inhibitor or miR-1298 inhibitor for 48 h (n=3 biological replicates). Differences were determined by tandem mass tag mass spectrometry analysis of protein lysates from the cells.

## DISCUSSION

The biological phenomenon that the endometrium acts as a biosensor of embryo developmental competency has been known for over a decade, however, the mechanism by which this is mediated has been elusive. We tested the hypothesis that differences in the cargo of EVs derived from conceptuses with different developmental potential are responsible for the biosensor capabilities of the endometrium. We have clearly demonstrated that the endometrium responds differently to embryos produced by assisted reproductive technologies and these differ depending on how the embryo is produced (Figure 4). These differences are also modified depending on the sex of the conceptus, extending this biosensor capability of the endometrium not only to developmental competency but to conceptus sex also. Following characterization of EVs from conceptuses during the pregnancy recognition period, differences in the miRNA cargo of EVs were determined. Treatment of endometrial epithelial cells with miRNA mimics and inhibitors for selected miRNAs altered the proteomic composition of the cells, proving for the first time that different miRNAs cargo from conceptus-derived EVs are responsible in part for the biosensor capability of the endometrium.

A major factor that can influence the response of the endometrium to the conceptus is via the actions of P4 on the endometrium which can alter the developmental trajectory of the conceptus (21). In order to exclude this as a confounding factor from our study we measured concentrations of P4 in the recipients. No differences were observed in the concentration of this key molecule (P4) that regulates the endometrial landscape. Therefore, any differences observed in the endometrial response are not due to the enhanced (22) or delayed (23, 24) P4 alterations the endometrium or ULF composition (4, 25). *In vitro* culture of the conceptus could lead to the identification of proteins that may be an artifact of the culture conditions in this study. In order to reduce this possibility, we have accurately characterized EVs according to current ISEV guidance (26). Moreover, previous work has sown that conceptus-derived proteins do have a functional effect on the endometrium (4, 11, 27), demonstrating that there are functional roles for these proteins identified in conceptus-conditioned medium (CCM). The differences observed are therefore due to differences in composition of the EVs themselves as no changes in size, concentration, or indeed markers were observed in EVs from different source conceptuses.

Our study has independently confirmed that the endometrium is a biosensor of developmental potential of the embryo mediated by EVs, as well as identified transcripts that are ART-specifically induced in the endometrium (Figure 4B). Moreover, we have shown that during the pregnancy recognition period the endometrium is also capable of deciphering between male and female conceptuses on Day 16 something not previously reported (28). Data in several mammalian species have shown embryos produced using different assisted reproductive technologies have different developmental potential and pregnancy outcomes (1, 5, 6). For the first time, our evidence demonstrates that these are potentially mediated via differences in embryo-maternal cross talk during the peri-implantation period of pregnancy. While we know this bi-lateral talk is necessary to support conceptus elongation (29, 30), and required to signal pregnancy recognition, these data support the notion that more nuanced interactions occur with continuous fine-tuning of the necessary secretion from the endometrium that is tailored to the needs (in terms of development) by the conceptus. Despite substantial differences in the transcriptome of the conceptus as it transitions from the blastocyst to an elongated conceptus (31), there has been limited data on the endometrial response to the sex of the conceptus (32). Our data show that there is endometrial sensitivity to male and female conceptus during the pregnancy recognition period.

By using an *in vitro* approach to transfect cells with selected mimics and inhibitors for selected miRNAs that are altered in conceptus-derived EV cargos, we have determined what key pathways in the endometrium that are altered via differences in EV cargo during pregnancy recognition. There are also likely differences in other cargo components. The majority of pathways altered are involved in the secretion and internal cell signaling. This may indicate that not only is the endometrium a biosensor, but that conceptus-derived EVs could potentially alter the endometrial secretome. One of the key functions of the endometrium is production of ULF to support growth and development of the conceptus prior to the formation of the placenta. It is possible that differences in EV cargo are produced to signal to the endometrium to modify the composition of ULF to help tailor to the specific developmental needs of the conceptus. We also provide additional evidence of the lack of robustness in terms of miRNA target prediction and that which can be demonstrated *in vitro*. In addition, we have determined new and novel targets of bovine miR128 and miR-1298 in a tissue specific manner, (i.e. endometrium), demonstrating the need for further tissue and cell specific target prediction models and in data bases such as the well curated mirGeneDB (33).

The phenomena of the biosensor capability of the endometrium in different mammalian species with different implantation strategies has been known for over a decade (5–7, 34, 35). However, the mechanism of action of how this is achieved has remained elusive. Our study clearly demonstrates that the response of the endometrium to embryos of different developmental potential (and sex) is mediated, in part, by different miRNA cargo in EVs derived from the conceptus. Specifically, we have shown that miR-128, and miR-1298 modify the endometrial response to embryos produced using ARTs compared to *in vivo* produced embryos. These data enhance our understanding of this biological phenomenon but also provide a road map for intervention to enhance developmental trajectory of the embryo and subsequent pregnancy outcome.

## MATERIALS AND METHODS

Unless otherwise stated all chemicals were sourced from Sigma. All animal work was carried out at the University of São Paulo Pirassununga with full ethical approval from the local ethical committee (approval number: CEUA 7920220219). All experiments were carried out in three experimental replicates i.e. three sets of experiments with all three experimental groups represented in each replicate. All recipient animals were from *Bos taurus indicus* (Nelore breed) and all three types of embryos were derived from *Bos taurus taurus* as described below. Animals were between 2 and 5 years old and were managed under good body conditions. Recipients were maintained in *Panicum maximum* pastures and had water *ad libitum* throughout the experiment. All animals enrolled in the study had their estrous cycles synchronized and blood samples were taken on 7 and 16 days and all animals were scanned for determining CL active by Doppler ultrasonography on Days 7, 16, and 20.

### In vivo embryo production

Holstein cows (approximately 2^nd^ parity and between 60 to 120 days post-partum) were scanned to ensure they were undergoing normal estrous cycles. On day -9 all animals received an intramuscular injection (i.m.) of 2 mg of estradiol benzoate (Sincrodiol, Ourofino Saúde Animal, Cravinhos, Brazil) along with an intravaginal progesterone-releasing device containing 1 g of P4 (Sincrogest, Ourofino Saúde Animal). On Day -5 each donor (n = 17 total) received serial decreasing i/m. injections of FSH (Folltropin, Vetoquinol, France) 12 hours apart (1&2: 400 IU, 3&4: 300 IU, 5&6: 200 IU, 7&8: 100 IU). After injection 8, the P4 device was removed and in the evening of day -1, all animals received an i.m. injection of 0.01 mg of GnRH (buserelin acetate, Sincroforte, Ourofino Saúde Animal). All animals were artificially inseminated on day 0 in the AM and PM using semen from the same bull and ejaculate as used for IVF embryo production. On day 7, the tail of the donors was swabbed with Ethanol and a 5 mL injection of lidocaine (company, place) was injected into the coccygeal space to administer a local anesthetic prior to flushing of the reproductive tract. Both the ipsilateral and contralateral uterine horns were flushed with a total volume of 1 liter of sterile PBS without any additives. This was passed through a collection cup fixed with a nylon filter (75 µm) and transported to the lab and the flush was searched under a stereomicroscope for embryos at the appropriate stage of development (late morula/early blastocyst). Identified structures were graded, and placed in IVF media (detailed below) for 2-3 hours prior to embryo transfer.

### In vitro embryo production

All *in vitro* embryos used for transfer into recipients were produced by Vitrogen (Cravinhos, SP). IVM was performed in TCM 199 supplemented with 10% FBS, 5.0 μg/mL luteinizing hormone (LH, Lutropia-V, Vetrepharm), 0.5 μg/mL follicle stimulating hormone (FSH, Folltropin-V, Vetrepharm), 0.2 mM pyruvate and 50 μg/mL gentamycin. After 22 h of IVM oocytes were *in vitro* fertilized (IVF) with thawed semen from the same bull and from the same ejaculate, prepared using a Percoll gradient (36). Fertilization (1 x 10^6^ sperm/mL) was performed in TALP medium, supplemented with 2 μM penicillamine, 1 μM hypotaurine, 250 μM epinephrine and 20 μg/mL heparin. After 18 h of fertilization the presumptive zygotes were mechanically stripped of cumulus cells by successive pipetting and transferred to *in vitro* culture (IVC) in SOF (synthetic oviduct fluid) medium supplemented with 2.5% FBS, 5.0 mg/mL BSA, 0.2 mM pyruvate and 50 μg/mL gentamycin. All IVM, IVF, and IVC procedures were performed in 100 μL drops (20 oocytes per drop), under mineral oil at 38.5°C in 5% CO_2_.

### Generation of cell lines and SCNT embryo production

To establishment of fibroblasts cell lines for cloned embryos, biopsies of subcutaneous tissue from the tail crease of one male and one female Holstein Friesian calf were collected at the Department of Dairy Cattle of University of São Paulo in the Fernando Costa campus. Biopsy fragments were cut into smaller pieces and digested in collagenase (1 mg/mL) for 3 hours at 37°C. Digested tissues were washed and centrifuged at 600 g for 10 min twice and cultured *in vitro* in α-MEM medium containing 20% FBS and 50 μg/mL antibiotic (penicillin and streptomycin). The medium was replaced every two days until the establishment of culture. Fibroblasts were transfected with the DNA constructs using a stable transducer with lentivirus vectors (MGH Vector Core, Boston, MA, USA) to express the genes of interest. The GFP (PalmGFP) constructs, provided by the researcher Charles L. Lai, Department of Neurology and Radiology, Massachusetts General Hospital Harvard Medical School (Lai et al., 2016). Briefly, lentiviral particles were produced by lipofection of 293FT cells (Invitrogen) with Lipofectamine 2000 (Invitrogen). The lipofection was incubated overnight (12-16h) and after this time the medium was completed replaced. Cells were seeded at a rate of 1 x 10^5^ cells per well on 6-well plates and 50 μL of the viral concentrate plus 8ng/mL polybrene (hexamethrin bromide) was added. Culture medium was replaced every 12 hours for 5 days. Cells were then frozen at -80°C in α-MEM medium containing 10% FBS, 10% DMSO and 50 μg/mL antibiotic and stored in liquid nitrogen.

### Flow cytometry and sorting of GFP-positive cells

After transfection, fibroblasts from both cell linages (male and female) were evaluated by flow cytometry to check the percentage of GFP-positive cells. As a negative control, non-transfected fibroblasts from both lineages were used. Briefly, the cells were disaggregated from the plates by enzymatic action (TrypLE Express^®^ - Thermo Fisher) per 5 min. After this time, the cells were centrifuged at 300 g for 10 min and the pellet was resuspended in 1 mL of fresh media. The cells were then stained with 10 μg/mL Hoechst 33342 for 15 min. Afterwards, the cells were evaluated separately using FACSCalibur flow cytometer (BD, Seattle, WA, USA). After sorting, the total of cells GFP-positive were recovered in a new 15 mL sterile tube, centrifuged at 300 g for 10 min and then were seeded *in vitro* for later use during the somatic cell nuclear transfer to generate cloned embryos positive for GFP.

### Somatic cell nuclear transfer

The production of the cloned blastocysts was performed according to (37, 38). Briefly, ovaries were collected from a commercial slaughterhouse and transported to the laboratory in physiological solution (NaCl 0.9%) at 35°C, supplemented with antibiotics (50 μg/mL penicillin and streptomycin). Follicles 3-8 mm in diameter were aspirated with the aid of 18 Gauge needle and 10 mL syringe and the recovered oocytes were selected using a stereomicroscope. Only those oocytes with a compact cumulus cell layer and homogenous cytoplasm (39) were used for *in vitro* maturation (IVM). Following 18 h of IVM, oocytes were denuded in 2% hyluronidase and those with the first polar body (1^st^ PB) were incubated in SOF medium containing 10 μg/mL Hoechst 33342 and 7.5 μg/mL cytochalasin B for 15 min. Micromanipulation was performed using an inverted microscope (Nikon Eclipse *Ti*-e, Tokyo, Japan) equipped with micromanipulators and microinjectors (Narishige, Tokyo, Japan). The 1^st^ PB along adjacent cytoplasm was removed by gentle aspiration using a glass pipette with 15μm of internal diameter (ES transferTip; Eppendorf, Hamburg, Germany). Before aspiration the oocytes were quickly exposed under UV filter (440 nm). Enucleated oocytes were reconstructed by the injection of a single fibroblast (control, male-derived GFP-positive, or female-derived GFP-Positive fibroblasts) into the perivitelline space of each oocyte. The oocyte and fibroblast resulting from the manipulation were fused in a fusion chamber (Eppendorf) filled with a fusion solution (0.28 M mannitol, 0.1 mM MgSO_4_, 0.5 mM HEPES, and 0.05% BSA in H_2_O) and subjected to alternating current (0.05 kV/cm for 5 sec) and direct current (1.75 kV/cm for 45μ sec). The presumptive zygotes were activated (26h after IVM) using 5 μM ionomycin in TCM HEPES 199 with 0.1% BSA, 0.2 mM sodium pyruvate and 50 μg/mL gentamicin for 5 minutes, followed by incubation in TCM-HEPES 199 with 3% BSA and in 2 mM 6-dimethylaminopurine (6-DMAP) in SOF for 3 h. After this period, the presumptive zygotes were transferred to the IVC as described above (100 μL drops ± 20 zygotes per drop).

### Embryo transfer

On Day 7 of the estrous cycle, all recipients were scanned by transrectal ultrasonography (Hz, and source) as described above to ensure the presence of an active CL. Blastocysts from all groups were graded and only those of grade I were individually loaded into transfer straws containing only 1 blastocyst each. Transfer straws were kept at 37°C in a portable incubator and randomly assigned to recipients.

### Assay for circulating P4 concentrations

Blood samples were collected on days 7 and 16 from the jugular vein for determination of plasma P4 concentrations. Samples were taken using a 10 mL vacuum tube containing heparin (BD Vacutainer, São Paulo, SP, Brazil). The samples were centrifuged at 3,600 g for 15 min at 4°C and the plasma was stored in the freezer at -20°C for subsequent measurements by a solid-phase RIA kit following the manufacture’s protocol (ImmuchemTM Double Antibody Progesterone Kit; Cat. 07e170105, MP Biomedicals, NY, USA). The intra-assay CV was 4.37%, and sensitivity was 0.077 ng/mL.

### Sample recovery

At day 16 of pregnancy, all reproductive tracts were processed for sample collection within 10 min following slaughter. Ipsi- and contra-lateral uterine horns were identified, and the reproductive tract was trimmed free of excess tissue to facilitate flushing of the uterine horns with laboratory grade PBS (pH 7.4). Ten mL of PBS were flushed through the ipsilateral and then contralateral uterine horns. The flush was collected in a sterile petri dish (100 mm) and the presence of an appropriately development conceptus was noted. Once the conceptus was identified, each were cultured individually in 3 mL of SOF media at 38.5°C for 6 h to allow collection of conceptus-derived EVs as previously described (4), as well as their contemporary blank. Following culture, the conceptus was removed from the culture media and the CCM as well as the contemporaneous blanks were processed for EV analysis as described below.

### Isolation and validation of EVs from conceptus-conditioned medium

Isolation of EVs (<200 nm) was carried at 4°C unless otherwise stated by standard differential centrifugation followed by ultracentrifugation as previously described (40) with minor modifications. Briefly, the CCM samples were centrifuged at 300 g for 10 min to remove cellular debris followed by 10 min at 2,000 g, and a 30 min spin at 16,500 g for 30 min to pellet large micro vesicles. The resulting supernatant was filtered through a 0.22 μm sterile syringe filter (Millipore), to remove particles greater than 200 nm, and centrifuged at 100,000 g for 70 min (Beckman Coulter, 70TI rotor, 4 mL polycarbonate tube, cat 355645). Following centrifugation, the supernatant was discarded, and the pellet was washed with excess PBS (pH 7.4). The washed suspension was centrifuged an additional time at 100,000 g for 70 min to pellet the EV fraction.

### NanoSight analysis

The resulting EV pellets were resuspended in 50 μL and 100 µL of PBS for characterization, respectively. For size and concentration determination, samples were kept at 4°C prior to analysis by Nano Tracking Analysis (NTA; NanoSight instrument NS300, Malvern, UK,). Videos (5 × 30 sec) were acquired after the manual introduction of EV samples (aliquot diluted 1:500 in PBS), at camera level 12 and 38.5°C, and by reference to 50, 100, and 150 nm calibration beads (Malvern) to verify accuracy. The EV samples were preserved at -80°C until further use. EV size and concentration between groups were compared on R (R version 3.6.3) using an ANOVA (function aov). Data were plotted using ggplot2 in R.

### Transmission electron microscopy

Transmission electron microscopy (TEM) was performed to visualize the size and morphology of the EVs contained in the CCM of the conceptuses (IV, IVF and NT). For this, the EVs from CCM (1 mL) from each group were isolated by ultracentrifugation (100,000g 70 min at 4°C) as previously described. The resulting pellets with the EVs were fixed with 2.5% glutaraldehyde, 0.1 M cacodylate and paraformaldehyde 4% (pH 7.2 to 7.4) for 2 h at RT. Afterwards, ultracentrifugation was performed and the EVs pellets were resuspended in ultrapure water (Milli-Q; Millipore Corporation, Merk, Burlington, MA, USA) and the samples were ultracentrifuged again. The isolated EVs were diluted in 50 µL of ultrapure water and placed on a copper grid coated with Pioloform® (Scientific Agar, Essex, UK) for 5 min. The grid was immediately placed in a drop of 2% aqueous uranyl acetate for 3 min. Excess solution was removed and the reading was performed in a transmission electronic microscope (FEI 200 kV model Tecnai20 emitter LAB6).

### Imaging and confirmation of GFP in conceptus and endometrial samples

Conceptus GFP-positive was visualized on the inverted phase contrast microscope (Nikon Eclipse TS100, Tokyo, Japan) at 10x magnification and quickly exposure time for the conceptus both prior to and after 6 h culture. Endometrial samples from GFP clone were also analyzed for GFP incorporation and endometrium counter-stained with DAPI (358 nm) and visualised using the epifluorescence microscope (Carl Zeiss, Thornwood, NY) with FITC filters (448 nm) and 400ms exposure times. PCR analysis and western blot for the GFP insert was carried out on endometrial samples as follows. The samples of CCM (1 mL) from conceptuses (IV, IVF and NT) were ultracentrifuged, as previously described in the manuscript, and the resulting pellet was evaluated for the presence of specific protein marker of EVs (endosome pathway marker): ALIX (1:750, sc49267, Santa Cruz Biotechnology, Dallas, TX, USA). As negative marker was used the proteins GRP78 (1:1000, sc-166490, Santa Cruz Biotechnology, Dallas, TX, USA). Briefly, after ultracentrifugation, the pellet containing the EVs was resuspended in 50 µL of RIPA buffer. Laemmli buffer and 2-mercaptoethanol were added to the samples (1:4) which were then denatured at 95°C for 5 min. For each Western Blot analysis, 12 µL of each sample was loaded onto 8% SDS-polyacrylamide gels and the run was performed at 100 V for 2 h. Proteins were transferred to a nitrocellulose membrane (#1620112 Bio-Rad) using Bio-Rad products and system for 2 h at 80V. Membrane blocking was performed with 5% BSA (Sigma-Aldrich) in Tris-buffered saline (1X - TBST; 100 mM NaCl, 0.1% Tween 20 and 50 mm Tris, pH 7.4) for 1 h. Afterwards, the membranes were incubated overnight at 4°C with the primary antibodies in 1X-TBST containing 1% BSA. The next day, the blots were washed three times for 5 min with 1X-TBST in a shaker and incubated with HRP-conjugated secondary antibody diluted in 1X-TBST (1:2000) for 1 h at RT. At the end of this period, the membranes were washed three more times with 1X - TBST and treated with Clarity™ Western ECL substrate (Bio-Rad) for protein visualization.

### EV RNA extraction and sequencing

RNA was extracted using the Qiagen miRNeasy kit (Qiagen, UK) and the Qiagen MinElute Cleanup Kit (Qiagen, UK) following the manufacturer’s instructions. A total of 700 μL of QIAzol (Qiagen, UK) were added directly to the EV samples. After 5 min incubation, 140 μL of chloroform was added, samples shaken vigorously for 15 sec, and then incubated for 3 min at RT. The tubes were centrifuged for 15 min at 12,000 g at 4°C to separate the phases. Three hundred and fifty μL of 70% ethanol was mixed with the aqueous phase by vortexing, applied to an RNeasy Mini Spin column and centrifuged for 15 sec at 8,000 g. The membrane was preserved at 4°C for >200nt RNA extraction. The flow through containing <200nt RNA was mixed with 450 μL of 100% ethanol and vortexed. The mix was added to an RNeasy MinElute Spin column and centrifuged for 15 sec at 8,000 g. The membrane was then washed with 500 μL of RPE buffer and centrifuged for 15 sec at 8,000 g and washed again with 500 μL of 80% ethanol. The ethanol was removed, and the membrane dried for 2 min at 8,000 g. The membrane was finished drying by a 5 min spin at 8,000 g with the lid open. Finally, the RNA was eluted in 14 μL of RNase-free water for 1 min at 8,000 g and the membrane was re-eluted with the flow through and snap frozen and stored at -80°C until further processing.

### Sequencing of small RNA species in EVs

All the RNA sequencing analysis was performed by Novogene (Cambridge, UK). Briefly, 3’ and 5’ adaptors were ligated to 3’ and 5’ end of small RNA, respectively. Then the first strand cDNA was synthesized after hybridization with reverse transcription primer. The double-stranded cDNA library was generated through PCR enrichment. After purification and size selection, libraries with insertions between 18∼40 bp were ready for sequencing on Illumina sequencing with SE50. The library was checked with Qubit and real-time PCR for quantification and bioanalyzer for size distribution detection. Quantified libraries will be pooled and sequenced on Illumina platforms, according to effective library concentration and data amount required.

### Analysis of the miRNA cargo of EV’s

Data were analysed through the LeedsOmics pipeline for small RNA sequencing. For the <200nt samples, adapter trimming and quality trimming and filtering were performed using Cutadapt. We used Bowtie to align the reads, featureCounts to quantify the reads and DESeq2 to identify the differentially-expressed-genes. The differentially expressed miRNAs were filtered for a fold change >1 or <-1 and a p-adjusted value <0.05.

### Target prediction

miRNA targets were predicted on TargetScan (targetscan.org; version 7.2). The list of predicted targets for each miRNA were retrieved and filtered for Cumulative weighted context++ score > or = -0.5, as the lowest the score, the most likely the miRNA targets the gene. Finally, the list of targets was compared between groups on Venny (https://bioinfogp.cnb.csic.es/tools/venny/; version 2.1). bta-miR-12034 could not be searched on TargetScan as it is not present on the miRNA database of the website.

### Endometrial RNA extraction and sequencing

RNA was extracted using the Qiagen mini miRNeasy kit (Qiagen, UK) and the Qiagen MinElute Cleanup Kit (Qiagen, UK) following the manufacturer’s instructions. A total of 700 μL of QIAzol (Qiagen, UK) were added directly to the tissue samples, and homogneised thoroughly using a TissueRupter (Qiagen, UK) for 15-90s. After 5 min incubation, 140 μL of chloroform was added, samples shaken vigorously for 15 s, and then incubated for 3 min at RT. The tubes were centrifuged for 15 min at 12,000 g at 4°C to separate the phases. Three hundred and fifty μL of 70% ethanol was mixed with the aqueous phase by vortexing, applied to an RNeasy Mini Spin column and centrifuged for 15 s at 8000 g.

The membranes binding the >200nt RNA were washed with 350μL RWT buffer for 15 seconds at 8000 g. Then the membranes were DNase treated (DNase I, Qiagen) for 15 min at room temperature. After this, they were washed with 300μL of RWT buffer for 15 s at 8000 g. Two more consecutive washes with 500 μL of RPE buffer were performed for 15 s at 8000 g before drying the tube for 1 min at 16,100 g with the lid open. The RNA was eluted with 50 μL of RNase-free water for 1min at 16 100 g. The membrane was re-eluted a second time with the flow through. At this point samples were quantified using a Nanodrop and then snap frozen and stored at -80°C until further processing.

### Sequencing of >200nt RNA species in endometrium

All the RNA sequencing analysis was performed by Novogene (Cambridge, UK). Firstly, ribosomal RNA was removed by rRNA removal kit, and rRNA free residue was cleaned up by ethanol precipitation. Subsequently, sequencing libraries were generated using the rRNA-depleted RNA and performing procedures as follow. Briefly, after fragmentation, the first strand cDNA was synthesized using random hexamer primers. Then the second strand cDNA was synthesized and dUTPs were replaced with dTTPs in the reaction buffer. The directional library was ready after end repair, A-tailing, adapter ligation, size selection, USER enzyme digestion, amplification, and purification. The library was checked with Qubit and real-time PCR for quantification and bioanalyzer for size distribution detection. Quantified libraries will be pooled and sequenced on Illumina platforms, according to effective library concentration and data amount required.

### Bioinformatics analysis endometrial RNA

The adapter sequences trimming, and quality filtering of the samples >200nt were performed by Cutadapt implemented in the Trimgalore pipeline (41). Sequencing reads of each sample were aligned using STAR (42) with standard parameters for alignment with the *Bos taurus* genome (Ensembl *Bos taurus* ARS-UCD1.2), and gene count was analyzed using featureCounts (43) implemented in the Rsubread package (44). Genes were considered expressed when they presented more than 5 counts in at least 70% of samples per group. Differential gene expression analysis was performed using the DESeq2 package (45), considering significance when the adjusted p values were less than 0.1 (Benjamini-Hochberg - “BH”) and the module of log2 foldchange was greater than 1. Additionally, we considered genes as differentially expressed if they were exclusively expressed in one group (at least 5 counts in all technical replicates), and not expressed in the other group (zero counts in all technical replicates) within comparison and using the function filterByExpr from edgeR package (46). Gene ontology analysis was performed using clusterProfiler (47) and pathways explored using Pathview (48). Data were visualized using R software, in which we primarily observed the classification, intensity, and difference in expression between groups. Exploratory data analysis was performed with principal component analysis using plotpca function from DESeq2 (44) and ggplot2 package (49), smearplots built with ggplot2 (49), heatmaps using Ward.D2 clusterization method from pheatmap package (50).

#### Functional analysis of selected miRNA on endometrial expression

##### miRNA mimic & inhibitor transfection

Primary bovine endometrial epithelial cells (n=3) isolated as previously described (11) were plated at 50,000 cells/well in a 24-well plate. The cells were incubated in a 5% CO_2_ environment at 38.5°C in 2 mL RPMI 1640 media (Gibco, UK) with 10% FBS (Gibco One-Shot FBS, UK) (charcoal stripped to deplete hormones), and 1% ABAM (Gibco, UK). Seventy-two hours after seeding, the cells were washed in PBS and the media was replenished with 450 μL RPMI media with EV-depleted charcoal-stripped FBS (Gibco One-Shot exosome-depleted FBS, UK) without antibiotics per well. The cells were transfected with the optimal concentration of 2 μL of lipofectamine 3000 (Invitrogen, UK) and 200nM mimic/inhibitor (Horizon Discovery, UK) per well. The experimental design comprised a control (50 μL OptiMEM only (Gibco, UK), a vehicle control (2 μL lipofectamine 3000 in 50 μL OptiMEM), a mimic and an inhibitor of each miRNA studied (miR-128 and miR-1298), a non-targeting mimic, and a non-targeting inhibitor. Briefly, per well, 2 μL of 100uM mimic/inhibitors were mixed 18 μL OptiMEM, then further diluted by taking 10 μL and adding a futher 15 μL OptiMEM (resulting concentration 4 μM). 2 μL of lipofectamine was mixed with 250 μL of OptiMEM medium and incubated for 5 min at RT. After the incubation, the lipofectamine solution (25 μL) was individually mixed with the targeting/non-targeting mimic/inhibitor solution (25 μL) and after a 20 min incubation at room temperature they were added to the corresponding wells. The cells were collected 48h after treatment by lifting with trypsin (0.025% for 5 min), neutralizing with medium and pelleted at 500 g for 5 min, re-suspended in PBS, re-pelleting at 500xg for 5 min, and cell pellet snap frozen prior to proteomic analysis.

##### Proteomic analysis

Cell pellets were placed on ice, and 40 μL RIPA buffer 1X (Merck, UK) with added cOmplete ULTRA protease inhibitors (Roche, UK), vortexed briefly, and kept on ice for 20 min. Samples were then centrifuged for 15 min at 14,000 g at 4°C and supernatant transferred to Eppendorf’s alongside a sample of lysis buffer for proteomics analysis. Tandem Mass Tag Labelling and High pH reversed-phase chromatography as carried out as previously described (51) by the University of Bristol core facility with minor modifications. These were the raw data files were processed and quantified using Proteome Discoverer software v2.1 (Thermo Scientific) and searched against the UniProt *Bos taurus* database (downloaded October 2021: 37512 entries) using the SEQUEST HT algorithm. The peptide precursor mass tolerance was set at 10ppm, and MS/MS tolerance was set at 0.6Da. The search criteria included oxidation of methionine (+15.995Da), acetylation of the protein N-terminus (+42.011Da) and Methionine loss plus acetylation of the protein N-terminus (-89.03Da) as variable modifications and carbamidomethylation of cysteine (+57.021Da) and the addition of the TMT mass tag (+229.163Da) to peptide N-termini and lysine as fixed modifications. Searches were performed with full tryptic digestion and a maximum of 2 missed cleavages were allowed. The reverse database search option was enabled and all data was filtered to satisfy false discovery rate (FDR) of 5%.

## ACKNOWLEDGEMENTS

We would like to acknowledge the help and assistance of students throughout the sample collection phase of this project. This work was supported by BBSRC grant number BB/R017522/1, QR-GCRF, and FAPESP (2016/22790-1, 2017/50438-3 and 2018/14137-1). We would like to acknowledge the assistance of the University of Leeds LeedsOmics facility, and the University of Bristol Proteomics core facility. We acknowledge Biorender in our production of components of Fig. 1. We would like to thank the staff and laboratory of flow cytometry from Hemocentro, Faculty of Medicine, University of São Paulo, Ribeirão Preto, SP, for the support performing the cytometry analyzes.

## DATA AVAILABILITY

The mass spectrometry proteomics data have been deposited to the ProteomeXchange Consortium via the PRIDE partner repository with the dataset identifier XXX. All RNA sequencing data are available via GEO database number XXX.

